# Long terminal repeats influence deletion propensity and genome instability in proviruses across major retroviruses

**DOI:** 10.64898/2026.10.12.758617

**Authors:** Sagnik Bhattacharjee, Koleta Michalek, Karin J Metzner

**Author notes:** Correspondence should be addressed to Karin J. Metzner.

## Abstract

A hallmark of retroviral infection is integration of viral genomes into host chromosomes to form proviruses. Most proviruses are defective, particularly in people with HIV receiving antiretroviral therapy, with large internal deletions as a dominant outcome. The contribution of retroviral long terminal repeats (LTRs) in the formation of large internal deletions remains unclear. We used dual-fluorophore reporters based on HIV-1, HIV-2, and HTLV-1 LTRs, integrated at AAVS1 by CRISPR/Cas9, to test whether LTRs influence proviral integrity in the absence of reverse transcriptase or integrase. Cerulean expression was driven by the LTR and mCherry by an independent human promoter. Double-positive (Cerulean⁺/mCherry⁺) cells were a proxy for intact vectors, whereas single mCherry⁺ cells indicated latent and/or defective proviruses.

Double-positive cells harboured intact vectors for all constructs. Non-inducible single mCherry⁺ populations were enriched for extensively deleted proviruses with HIV-1, HIV-2, or HTLV-1 LTRs, whereas the LTR-deficient reporter yielded few single mCherry⁺ cells, all with intact proviruses. The presence of LTRs was significantly associated with deleted genomes. HIV-1 and HTLV-1 showed higher observed deletion rates than HIV-2, but this was not statistically significant owing to limited sequence numbers. Single Cerulean⁺ populations across all constructs showed recurrent deletions affecting the mCherry cassette and upstream promoter, with short direct repeats at deletion junctions. These likely reflect microhomology-mediated end joining as a contributing repair mechanism during CRISPR/Cas9-mediated integration, in which LTRs may not directly contribute. Together, these data identify retroviral LTRs as contributors to the formation of large internal deletions in proviruses.

**Importance:** Retroviruses such as human immunodeficiency virus type 1 (HIV-1), human immunodeficiency virus type 2 (HIV-2), and human T-lymphotropic virus type 1 (HTLV-1) cause lifelong infections that can lead to AIDS, leukaemia, and other severe diseases, respectively. After infection, these viruses convert their RNA into DNA and integrate it into the host genome, creating proviruses that can persist for years. Most proviruses are defective and often lack large parts of their genome, particularly in people with HIV receiving antiretroviral therapy, but the viral features that drive this instability have remained incompletely understood. Here we show that the long terminal repeats, which are specialized DNA sequences at the ends of the viral genome, influence the formation of defective proviruses. These insights are expected to improve our understanding of defective provirus landscapes in infected individuals and can inform the design and interpretation of retrovirus-based vectors in research and therapy.

## Introduction

Retroviruses cause chronic, often life-threatening diseases in humans. Human immunodeficiency virus type 1 (HIV-1) and human immunodeficiency virus type 2 (HIV-2) cause acquired immunodeficiency syndrome (AIDS), while human T-lymphotropic virus type 1 (HTLV-1) is associated with adult T-cell leukaemia/lymphoma (ATL) and inflammatory and neurological diseases (1, 2). After infection, retroviruses reverse transcribe their RNA genome into DNA and integrate the resulting DNA into host chromosomes, generating proviruses that can persist for years and form long-lived viral reservoirs that represent a major barrier to cure (1, 3–5).

In people with HIV-1 on antiretroviral therapy (ART), the great majority of integrated proviruses are defective, with up to 93–98% in CD4⁺ T cells carrying defects (6–8). Notably, defective proviruses arise early during infection: in a single-round in vitro HIV-1 infection model, 41% of proviral sequences were defective, indicating that defective proviruses can be generated at high frequency during the initial rounds of reverse transcription and integration (6). Similar observations have been made in ART-treated HIV-2 and simian immunodeficiency virus (SIV) infections, and a substantial fraction of the HTLV-1 proviral reservoir in ATL is also defective (9–13). Across these retroviruses, the most frequent defects are large internal deletions (LID), which vary in size and position and can remove internal segments >6 kb (6, 7, 9, 14). In HTLV-1, defective proviruses often fall into two categories, one with internal deletions and another with preferential deletion of the 5′ long terminal repeat (LTR) and upstream region (10). Human endogenous retroviruses (HERVs), remnants of ancient retroviral infections embedded in the human genome, likewise persist predominantly as deleted or mutated proviruses (15). Together, these observations highlight large internal deletions as a recurrent and prominent feature of retroviral proviral landscapes.

LID is often attributed to errors during reverse transcription, supported by short direct repeats at some deletion junctions and by immune- or treatment-driven selection, but such repeats are detected in only a subset of cases and the mechanisms underlying most LID remain incompletely understood (6, 8, 16–19). We previously showed that LID occur in HIV-1-based vectors and in HIV-1 sequences integrated into the human genome via CRISPR/Cas9 in the absence of viral reverse transcriptase and integrase, implicating host DNA repair pathways in the formation of defective proviruses (20, 21). In our recent study, we used a targeted CRISPR/Cas9 screen with the HIV-1-derived dual-fluorophore vector LTatC[M]L, which discriminates between double-positive Cerulean⁺/mCherry⁺ (DP) cells with intact integrated vectors and single mCherry⁺ cells with vectors carrying large internal deletions, to identify DNA damage response proteins that contribute to LID formation (22) . Clonal analyses in our previous studies showed that nearly all non-inducible single mCherry⁺ cells contained extensive deletions in the HIV-1 Tat and/or Cerulean region, whereas double-positive cells retained intact vectors (20, 21).

Building on this reporter, we generated a modified vector, LTatCL[M], in which the Cerulean cassette is positioned between two HIV-1 LTRs while the mCherry cassette remains outside the LTRs under a cellular promoter (20). The construct is integrated using CRISPR/Cas9 into the Adeno-associated virus integration site 1 (AAVS1) genomic safe harbour locus, with homology arms flanking the vector and unnecessary viral elements and packaging signals removed.

LTRs are distinctive elements that flank retroviral genomes and harbour regulatory and structural features that influence reverse transcription, integration and gene expression (23, 24). Given their conserved presence across retroviruses, we hypothesised that LTRs contribute to the formation of deletions in proviruses and that LTRs from different human retroviruses may shape the propensity of proviruses to acquire large internal deletions, even in the absence of viral enzymes. Here, we adapt the LTatCL[M] system to incorporate LTRs from HIV-1, HIV-2 or HTLV-1, as well as an LTR-deficient control, and use CRISPR/Cas9-mediated integration into the AAVS1 locus to dissect whether viral LTR architecture influences the formation of defective proviruses. In these systems, HIV-1 Tat is the only viral protein expressed and can transactivate heterologous retroviral LTRs, allowing comparison of different LTRs while keeping the transactivator constant (25–29). Our findings reveal that retroviral LTR architecture contributes to proviral deletion propensity and provide a controlled framework to study how conserved viral sequence elements shape the proviral landscape and genome stability alongside host and other viral factors.

## Results

### Retroviral long terminal repeats promote the formation of large internal deletions in integrated proviruses

We performed targeted integration of four model vectors, HIV-1 LTatCL[M], HIV-2 LTatCL[M], HTLV-1 LTatCL[M], and the LTR-deficient control ΔLTatCΔL[M] (Figure 1A), into the AAVS1 safe harbour locus in K562 cells by CRISPR/Cas9-mediated homologous recombination using the PX458_AAVS1 gRNA plasmid. Flow cytometric analyses were performed on days 6, 13 and 20 after nucleofection and live cells were gated using Zombie NIR staining before analysis. From day 13 onwards, all three retroviral LTR-containing vectors consistently yielded three clearly separated populations: double-positive Cerulean⁺/mCherry⁺ (DP), single mCherry⁺ and single Cerulean⁺ cells. In contrast, the LTR-deficient vector predominantly generated a strong DP population with high reporter expression driven by the human constitutive promoter, together with only a minor single mCherry⁺ population and a markedly lower single mCherry⁺/DP ratio (Figure 1B). This indicates that the presence of retroviral LTRs strongly favours the emergence of single mCherry⁺ cells, which we use as a proxy for defective or non-inducible proviruses carrying large internal deletions (LID) and/or exhibiting stable latency.

**Figure 1.**
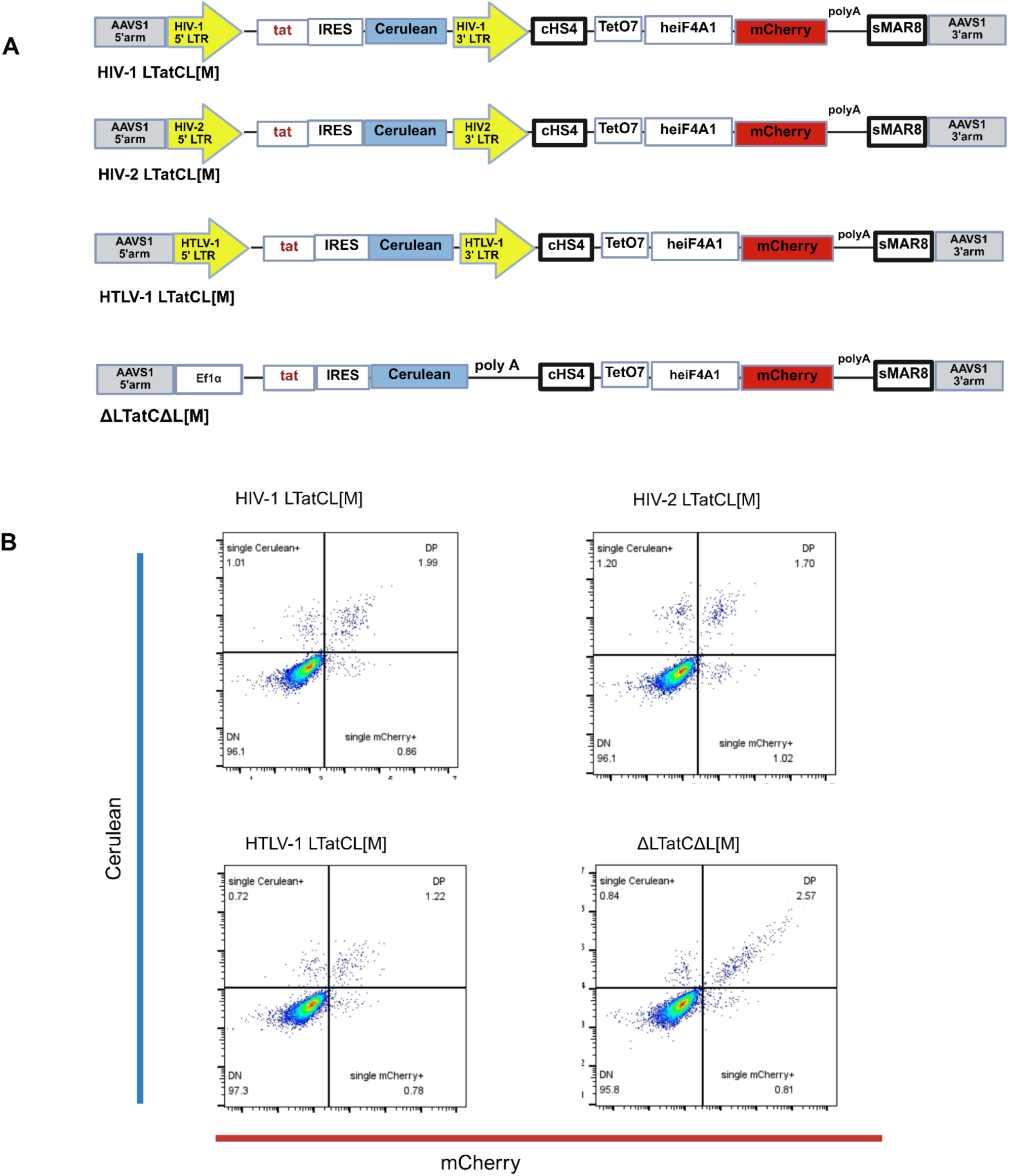
Retroviral long terminal repeats promote the formation of large internal deletions in integrated proviruses. A) Schematic representation of the four LTatCL[M] constructs. From top to bottom: HIV-1 LTatCL[M], HIV-2 LTatCL[M], HTLV-1 LTatCL[M], and ΔLTatCΔL[M]. Schematics were created with BioRender. B) Representative flow cytometry plots at day 20 after nucleofection of K562 cells with the four vectors, showing Cerulean versus mCherry fluorescence. Quadrants indicate double-positive (DP), single Cerulean⁺, single mCherry⁺, and double-negative (DN) populations. Data were analysed in FlowJo; plots were arranged and labelled in BioRender.

### Enrichment of single Cerulean⁺ and single mCherry⁺ subpopulations for analysis of large internal deletions

Building on these observations, we nucleofected the four vectors into K562 cells at large scale and performed an initial FACS to isolate defined subpopulations. The overall experimental workflow is shown in Figure 2A. For the three retroviral LTR-containing vectors (HIV-1-, HIV-2- and HTLV-1 LTatCL[M]), a prominent single mCherry⁺ population was already evident relative to DP cells (Figure 2B, and Figures S1–S2), whereas the LTR-deficient ΔLTatCΔL[M] vector showed a dominant DP population and only a minor single mCherry⁺ subset (Figure S3). Sorted populations were expanded separately and subsequently analysed by flow cytometry once each fraction had stabilised into a characteristic phenotype (Figure 2B and Figures S1–S3). Live cells were gated using Zombie NIR staining, as above. The populations were then subjected to a second FACS to collect highly purified subpopulations (Figure 2B and Figures S1–S3).

**Figure 2.**
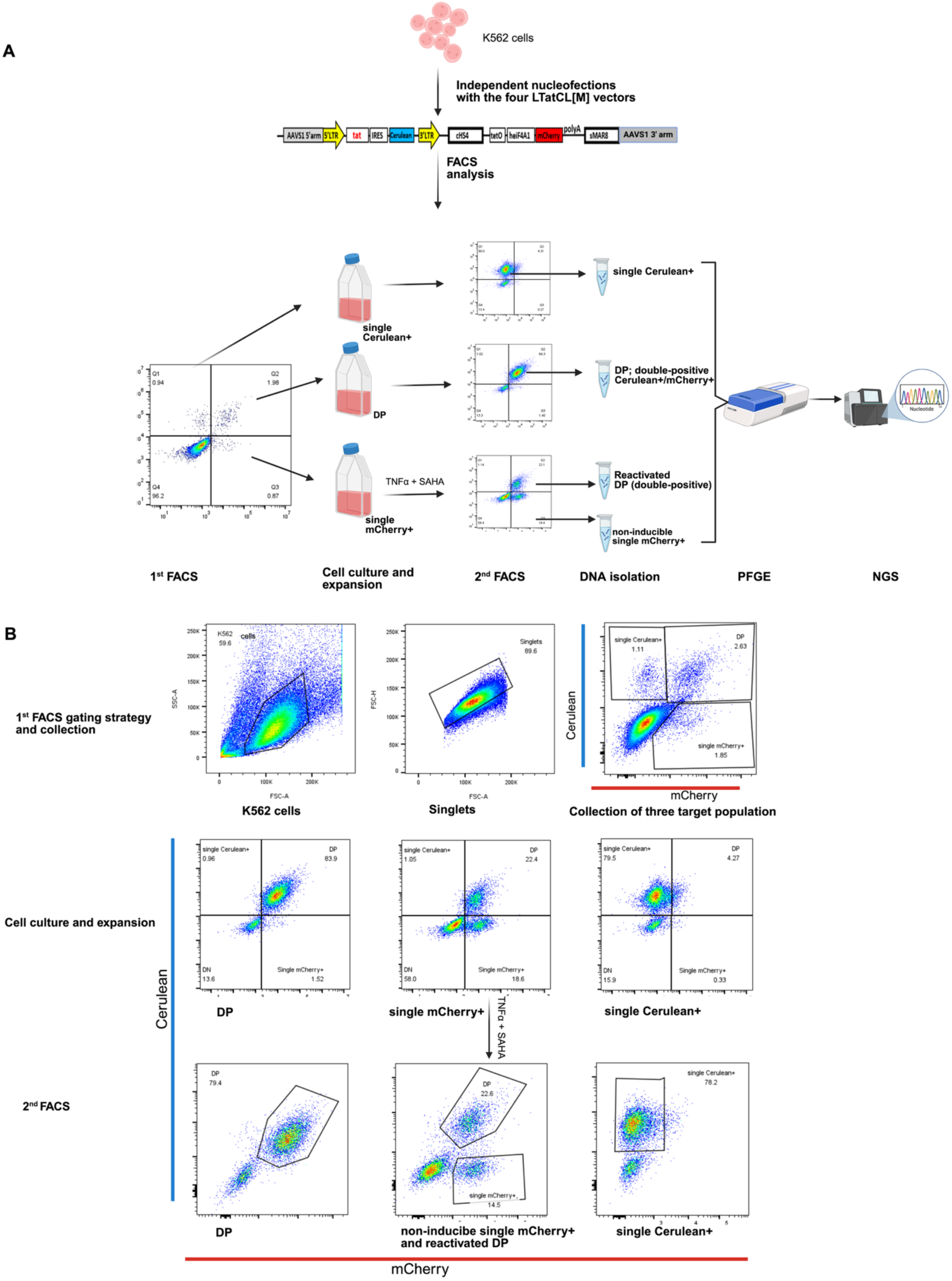
Enrichment of defined Cerulean⁺ and/or mCherry⁺ subpopulations for analysis of large internal deletions. A) Schematic workflow of the experimental strategy. Created with BioRender. B) Representative flow cytometry plots for HIV-1 LTatCL[M]. Top row: gating strategy used at the first sort (day 6 after nucleofection) to collect double-positive (DP), single Cerulean⁺, and single mCherry⁺ populations. Middle row: analysis of the expanded populations at day 16. Bottom row: gating at the second sort (day 18) to collect DP from the DP culture, non-inducible single mCherry⁺ and reactivated DP from LRA-treated single mCherry⁺ cultures, and single Cerulean⁺ from single Cerulean⁺ cultures. Flow cytometry data from both analytical runs and sorting (CytoFLEX and BD FACSAria) were reanalysed in FlowJo to apply consistent quadrants, axis orientation, and labelling across experiments; plots were arranged and refined in BioRender without altering underlying event distributions.

DP-sorted cells remained predominantly DP for all vectors. In contrast, single mCherry⁺-sorted cells from HIV-1 LTatCL[M] and HIV-2 LTatCL[M] contained both DP and single mCherry⁺ cells after expansion. As single mCherry⁺ cells can also indicate latent integrated proviruses, some cells may have auto-reactivated to the DP state, thereby contributing to the DP population within the single mCherry⁺-sorted fraction. A small contribution from sorting imprecision at the initial sort cannot be excluded.

To distinguish inducible latent from non-inducible defective single mCherry⁺ cells, expanded single mCherry⁺-sorted populations from each vector were treated for 24 h with latency reversal agents (LRAs) tumour necrosis factor alpha (TNF-α) and suberoylanilide hydroxamic acid (SAHA) prior to the second sort and then re-sorted into DP- and single mCherry⁺ fractions. For HIV-1 LTatCL[M] and HIV-2 LTatCL[M], reactivation produced only modest increases in the DP fraction, in line with our earlier observations that the majority of single mCherry⁺ cells remain non-inducible. (21) HTLV-1 LTatCL[M] behaved similarly overall but showed an even more stably predominant single mCherry⁺ phenotype before and after reactivation, whereas ΔLTatCΔL[M] yielded only a small single mCherry⁺ population with the lowest frequency of single mCherry⁺ cells among all vectors. A small single Cerulean⁺ population was observed and stably expanded for all four vectors (Figure 2B, Figure S1–S3). This previously underappreciated population is of interest, as DP cells serve as a proxy for intact proviruses and non-inducible single mCherry⁺ cells for integrated vectors with LID, whereas single Cerulean⁺ cells likely reflect defects or deletions within the mCherry cassette.

These sorted DP-, non-inducible single mCherry⁺-, reactivated DP- and single Cerulean⁺ populations formed the basis for subsequent sequencing based mapping of large internal deletions. The overall experimental workflow and FACS plots with gating strategies are shown in Figure 2 and Figures S1–S3.

### Enrichment of LTR-dependent deleted genomes in non-inducible single mCherry⁺ populations

Genomic DNA from sorted DP, reactivated DP, non-inducible single mCherry⁺ and single Cerulean⁺ populations of all four vectors was enriched for high molecular-weight fragments containing integrated vectors by pulsed-field gel-electrophoresis (PFGE), thereby depleting episomal and short DNA species. Purified DNA was then used as template for long-range PCR spanning the Cerulean cassette (from tat to the start of mCherry) in DP, single mCherry⁺ and reactivated DP populations, and for PCR of the mCherry cassette (from the middle of Cerulean to the start of the 3′ AAVS1 arm) in single Cerulean⁺ populations. For each condition, multiple independent PCRs were performed, and reactions yielding discrete bands were subjected to next-generation sequencing.

For all four vectors, DP populations yielded predominantly single full-length amplicons on agarose gels, and deep sequencing confirmed intact Cerulean and mCherry cassettes in all analysed HIV-1 LTatCL[M], HIV-2 LTatCL[M], HTLV-1 LTatCL[M], and ΔLTatCΔL[M] DP amplicons (Figure S5A). In HIV-1 LTatCL[M] reactivated DP cells derived from single mCherry⁺-sorted populations, 5 of 6 sequenced amplicons were fully intact. One amplicon (HIV-1 rDP13) carried an internal deletion confined to the downstream CHS4 insulator/tetO7/EF1α region, leaving both Cerulean and mCherry cassettes intact and preserving DP expression, consistent with residual EF1α activity and/or upstream LTR-driven transcription (Figure 4B and Figure S5B). In HTLV-1 LTatCL[M] reactivated DP cells, 3 of 4 amplicons were intact, whereas one (HTLV-1 rDP01) harboured a large deletion extending from the Cerulean region into the EF1α promoter yet still scored DP by flow cytometry, consistent with strongly reduced but not completely absent Cerulean coverage and suggesting that low-level Cerulean expression from residual promoter fragments or upstream regulatory elements can suffice for DP classification in some cases (Figure 4B and Figure S5B). Reactivated DP amplicons from HIV-2 LTatCL[M] and ΔLTatCΔL[M] were all intact (Figure S5B).

In contrast, non-inducible single mCherry⁺ populations were strongly enriched for deleted genomes. Among non-inducible HIV-1 LTatCL[M] single mCherry⁺ amplicons, 7 of 11 carried LID spanning most of the Cerulean cassette and flanking sequences, whereas all five ΔLTatCΔL[M] single mCherry⁺ amplicons were intact. For HIV-2 LTatCL[M], 5 of 14 single mCherry⁺ amplicons were defective, and for HTLV-1 LTatCL[M], 5 of 7 were defective with LID. Representative coverage plots for five such single mCherry⁺ amplicons per vector (including one intact example) are shown in Figure 3. Mapping of deletion breakpoints and exact sequences at such junctions for all deleted non-inducible single mCherry^+^ amplicons is shown in Figure 4A. Deletions in such single mCherry⁺ amplicons frequently extended from Cerulean into downstream regions and, in some cases, into the mCherry promoter and coding sequence, indicating that truncated promoter elements and/or upstream LTRs can sustain sufficient (potentially truncated) mCherry expression to generate a single mCherry⁺ phenotype even when large parts of the cassette are missing. The observed fraction of defective single mCherry⁺ genomes was higher for HIV-1 and HTLV-1 LTatCL[M], lower for HIV-2 LTatCL[M], and absent for the LTR-deficient ΔLTatCΔL[M] vector. Across vectors, the presence of retroviral LTRs was significantly associated with LID, as assessed by Fisher’s exact test (*P* = 0.0498). Microhomologies or short repeats were not detected at deletion junctions in single mCherry⁺ amplicons. These findings support a role for retroviral LTR architecture in determining deletion propensity in integrated proviruses.

**Figure 3.**
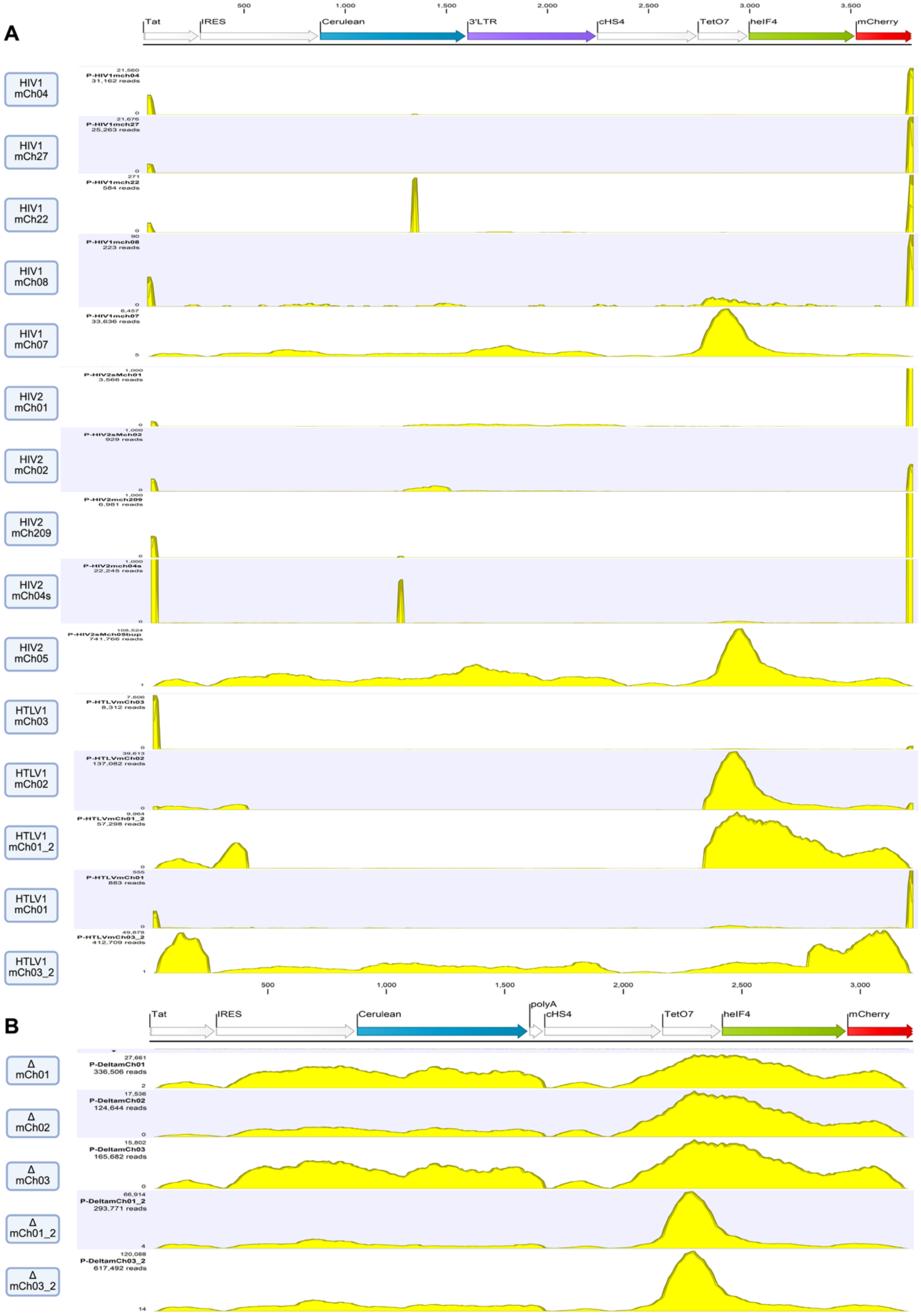
Enrichment of LTR-dependent deleted genomes in non-inducible single mCherry^+^ populations. A) Representative coverage plots of five amplicon sequences per vector showing deletions within the Cerulean cassette in non-inducible single mCherry⁺ populations for HIV-1 LTatCL[M], HIV-2 LTatCL[M], and HTLV-1 LTatCL[M]. At the top, a schematic of the corresponding entire reference region is shown. Yellow peaks indicate regions with read coverage (intact sequence), whereas flat gaps between peaks indicate internal deletions. In each set, the fifth amplicon illustrates an intact proviral segment without deletions. B) The LTR-deficient ΔLTatCΔL[M] construct is shown with its corresponding reference schematic and five representative amplicon sequences, all of which are intact and lack internal deletions. Individual amplicon identifiers (left) correspond to specific PCR products and sequencing reactions and match the sequence names used for data deposition. Coverage plots were exported from CLC Genomics Workbench after mapping reads to the corresponding reference constructs and assembled in BioRender.

**Figure 4:**
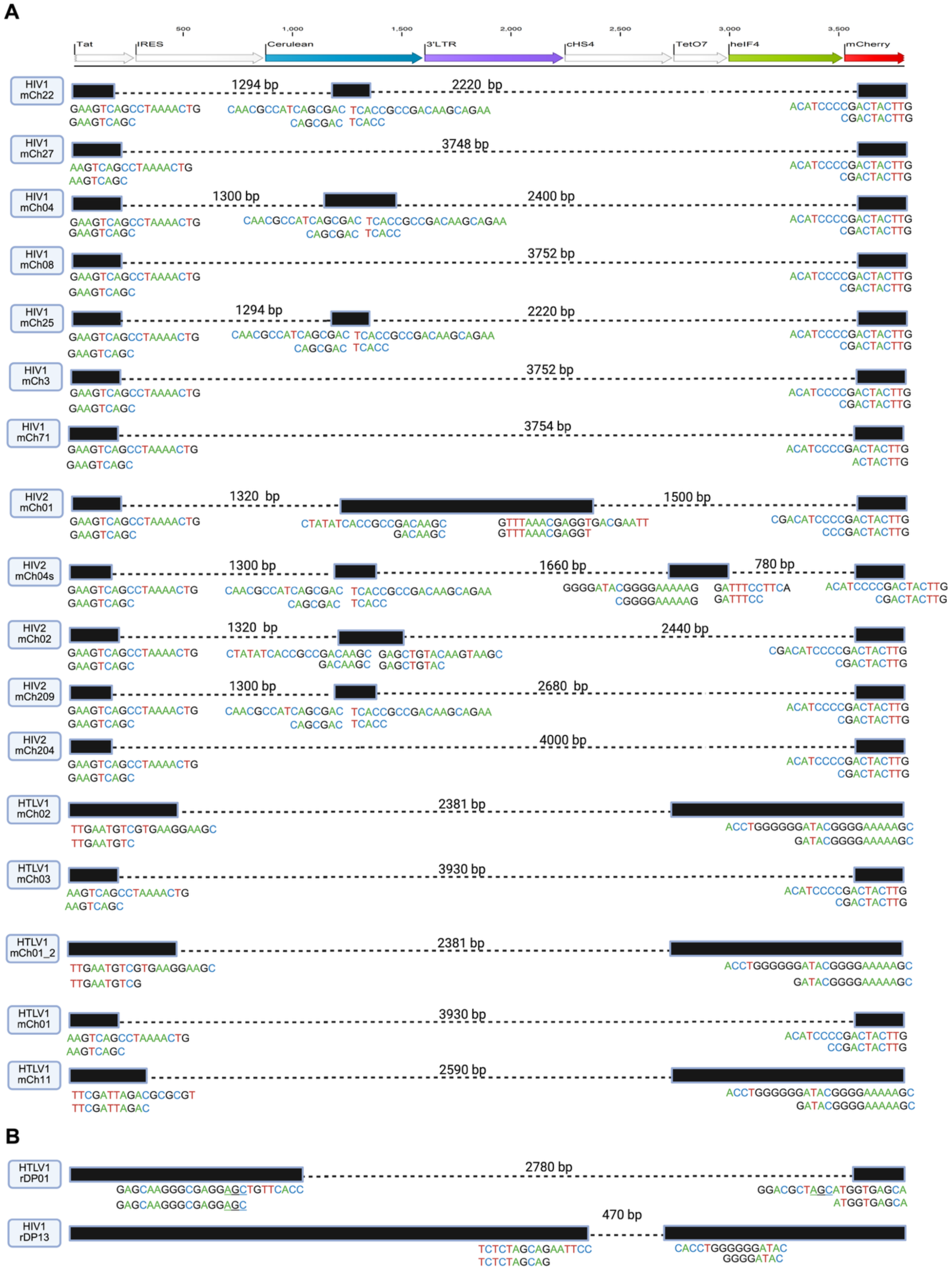
Mapping of deletions within the mCherry cassette in non-inducible single mCherry^+^ populations and exceptional reactivated double-positive populations from LTR-containing vectors. A) Schematic mapping of deletions in non-inducible single mCherry^+^ populations. B) Schematic mapping of deletions in two reactivated double-positive (rDP) amplicons from LTR containing vector, representing exceptions to the predominantly intact rDP genomes. Black bars indicate intact regions of each amplicon, and dotted lines indicate deleted regions. For each deletion, sequences flanking the breakpoint are shown beneath the schematic. The upper sequence corresponds to the intact reference region, and the lower sequence corresponds to the observed amplicon sequence; the junction between the flanking sequences marks the deletion breakpoint. At the top, a schematic of the corresponding enitre reference region is shown. Individual amplicon identifiers (left) correspond to specific PCR products and sequencing reactions and match the sequence names used for data deposition. Created with BioRender.

Segments with complete loss of coverage (0 reads) or with coverage reduced by more than two orders of magnitude relative to neighbouring regions were classified as true deletions. Occasional residual reads within such regions likely reflect sequencing noise or contributions from minor amplicon species within the same reaction. Coverage across the repetitive CHS4 insulator regions was generally reduced and noisy for all vectors, consistent with low mappability of these sequences, and was therefore interpreted with caution. Although the observed frequency of defective proviruses was higher in HIV-1 and HTLV-1 LTatCL[M] than in HIV-2 LTatCL[M], the limited number of sequenced amplicons precludes a definitive comparison among the individual LTR-containing vectors and warrants further investigation.

### Large internal deletions and microhomology signatures in single Cerulean⁺ populations

Analysis of single Cerulean⁺ populations revealed a deletion pattern present in all LTR containing and LTR deficient vector. For each vector, four or five single Cerulean⁺ amplicons were sequenced, and all contained deletions within the mCherry cassette, including those from ΔLTatCΔL[M] (Figures 5 and S4). In many cases, deletions extended from the downstream CHS4 insulator through the EF1α promoter and mCherry coding region, consistent with the absence of mCherry expression. Inspection of deletion junctions in single Cerulean⁺ amplicons frequently revealed short direct repeats shared between the upstream and downstream breakpoints, including 3–8-bp motifs such as GGTG or CCGGA (Figure 5). Coverage plots for such single Cerulean^+^ amplicons per vector are shown in Figure S4. Mapping of deletion breakpoints and the corresponding junction sequences for all deleted single Cerulean⁺ amplicons is shown in Figure 5.

**Figure 5.**
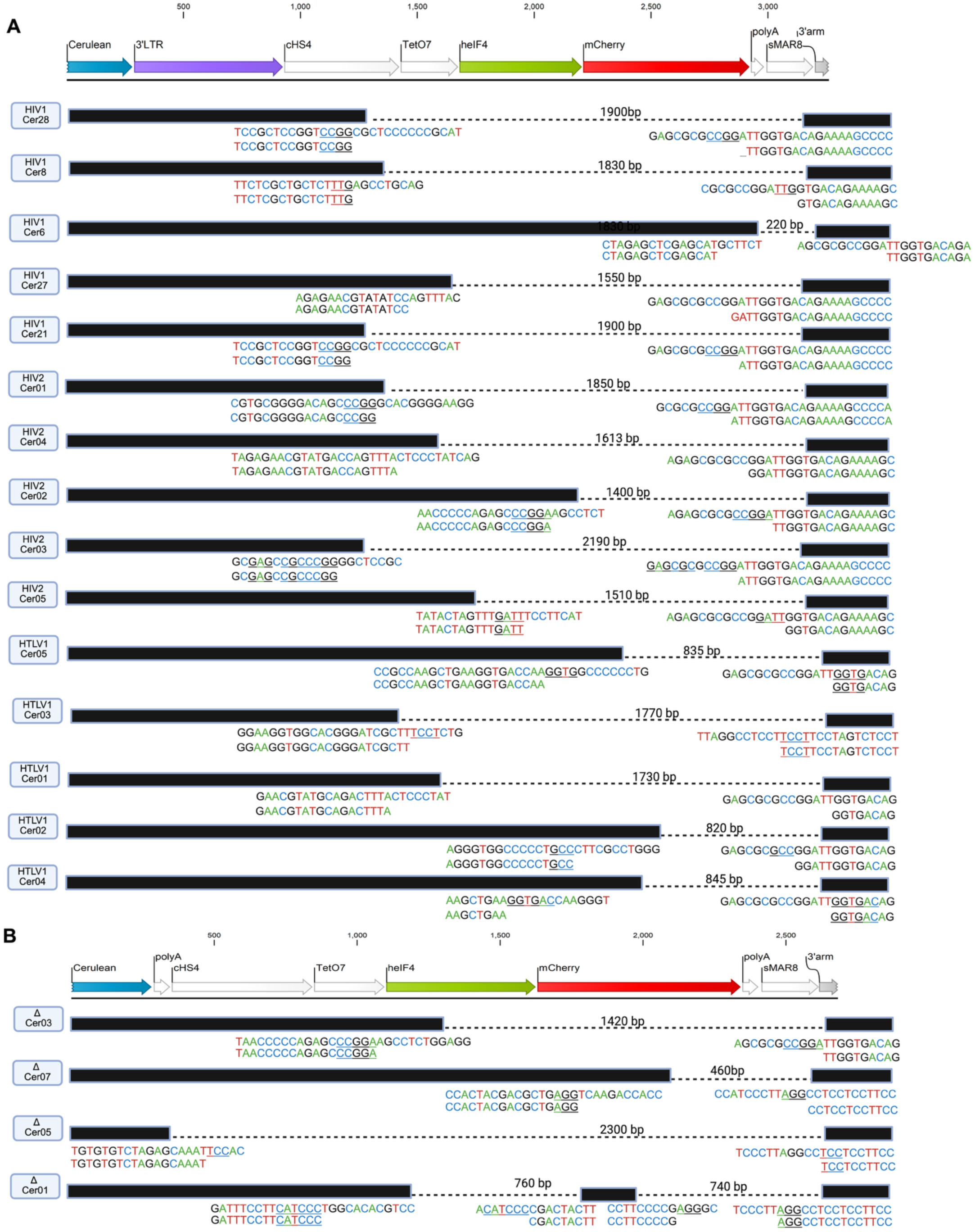
Large internal deletions and microhomology signatures in single Cerulean⁺ populations. Schematic plots of all amplicon sequences per vector showing deletions within the mCherry cassette in single Cerulean⁺ populations for (A) HIV-1 LTatCL[M], HIV-2 LTatCL[M], and HTLV-1 LTatCL[M], and (B) the LTR-deficient ΔLTatCΔL[M] construct. Black bars indicate intact regions of each amplicon, and dotted lines indicate deleted regions. For each deletion, sequences flanking the breakpoint are shown beneath the schematic. The upper sequence corresponds to the intact reference region, and the lower sequence corresponds to the observed amplicon sequence; the junction between the flanking sequences marks the deletion breakpoint. Microhomologous repeats at the junction are underlined. At the top, a schematic of the corresponding entire reference region is shown. Individual amplicon identifiers (left) correspond to specific PCR products and sequencing reactions and match the sequence names used for data deposition. Created with BioRender.

These junctional microhomologies are consistent with a potential contribution of microhomology-mediated end joining (MMEJ) to deletion formation. Because the same pattern was observed across all vectors, including the LTR-deficient construct, it is unlikely to be explained solely by the presence or identity of retroviral LTRs.

## Discussion

Our dual-fluorophore CRISPR/Cas9 integration system shows that proviruses in double-positive populations are predominantly intact, whereas non-inducible single mCherry⁺ and single Cerulean⁺ populations are strongly enriched for large internal deletions (LID) affecting the Cerulean and mCherry cassettes, respectively. Following reactivation with LRAs, single mCherry⁺ populations remained largely unchanged across the LTR-containing vectors, with limited conversion to double-positive cells; this was particularly evident for HTLV-1 LTatCL[M], in which single mCherry⁺ cells remained predominant before and after reactivation. Thus, reactivated double-positive populations were predominantly intact, whereas non-inducible single mCherry⁺ populations were enriched for deleted vectors.

The LTR-deficient ΔLTatCΔL[M] vector showed the lowest frequency of single mCherry⁺ cells, and all amplicons recovered from its non-inducible single mCherry⁺ population were intact by gel electrophoresis and sequencing. In contrast, LID occurred frequently in corresponding populations from vectors containing retroviral LTRs, and the association between LTR presence and deletion occurrence was statistically significant. These findings indicate that, in this system, two retroviral LTRs promote LID formation. Although intact and deleted vector proportions varied among HIV-1, HIV-2, and HTLV-1 LTR constructs, these differences were not statistically significant owing to the limited number of analysed sequences. Thus, individual LTRs cannot be concluded to generate different deletion frequencies, although LTR sequence and architecture may modulate LID frequency without being the sole determinants of deletion formation. Interestingly, retroviral proviral landscapes can contain highly deleted genomes retaining only LTR-derived sequences, including solo LTR forms described in HIV-1 reservoirs and endogenous retroviruses, raising the possibility that retroviral terminal structures contribute more broadly to defective proviral genomes (30).

These findings extend previous work implicating host DNA damage response factors in the generation of large internal deletions in integrated HIV-1 sequences (22) and suggest that conserved retroviral terminal structures contribute to shaping the proviral landscape alongside viral enzymes and host repair pathways. Future studies should determine whether specific LTR features influence host DNA damage response pathways and how this is modified by other viral proteins, integration-site context, and cell type.

Despite the strong enrichment for deleted genomes, a subset of non-inducible single mCherry⁺ clones retained intact vectors in both LTR-deficient and LTR-containing vectors, consistent with transcriptional silencing rather than structural defects. HIV-1 latency is governed by epigenetic regulation at the LTR, including nucleosome positioning and repressive complexes such as HDACs and YY1 (31). Because LRAs such as TNF-α and HDAC inhibitors only partly reactivate the latent reservoir (32, 33), a similar mechanism may underlie the intact but transcriptionally inactive HIV-2 LTatCL[M] vectors. HIV-2 LTR-driven transcription is weaker and less responsive to activation signals than HIV-1, and HIV-2-based constructs may therefore be more prone to transcriptional silencing and reduced LRA reactivation (34, 35). This may explain the higher frequency of intact but non-inducible single mCherry⁺ clones in the HIV-2 context, although the difference among retroviral LTR-containing vectors was not statistically significant.

Single Cerulean⁺ amplicons showed recurrent deletions spanning the CHS4 insulator, EF1α promoter, and mCherry cassette across all vectors, including the LTR-deficient construct, with frequent 3–8-bp direct repeats at deletion junctions. These microhomologies are consistent with MMEJ or related error-prone end joining following CRISPR/Cas9-induced double-strand breaks at AAVS1, particularly across repetitive sequences such as the CHS4 insulator. Cas9-induced large deletions are often microhomology-associated, and POLQ-dependent MMEJ can compete with HDR; inhibiting MMEJ can suppress such events while enhancing HDR in the presence of a donor template (36–38). These deletions may therefore represent occasional MMEJ-like outcomes alongside successful HDR-mediated AAVS1 targeting, reflecting the local repair environment rather than a direct LTR effect. This interpretation warrants further investigation and highlights the need to consider large microhomology-associated deletions in CRISPR-based editing strategies (39) .

This study is limited to retroviral vectors and a transformed cell line in the absence of most viral proteins; the quantitative contribution of LTR architecture may therefore differ in primary cells, at diverse integration sites, and during complete viral replication cycles. Nevertheless, this work provides an initial framework to dissect how retroviral terminal architecture contributes to defective proviruses. LTR variants selectively removing U3 or R/U5 segments, together with analyses in primary cells and patient-derived proviruses, should help identify the relevant LTR subregions and cellular contexts.

## Materials and Methods

### Cell culture

K562 cells (ATCC CCL-243) were cultured in RPMI-1640 (Sigma-Aldrich, R8758) supplemented with 10% (v/v) fetal bovine serum (FBS; Thermo Fisher Scientific, A5256701), 100 U/ml penicillin (Gibco, 15140122), and 100 μg/ml streptomycin (Gibco, 15140122). MT-2 cells were cultured similarly as K562, but under BSL-3 conditions. All cell lines were passaged every 2–3 days at a 1:10 split ratio and incubated at 37°C in 5% CO₂.

### Plasmid constructs

The LTatCL[M] model vector contains the HIV-1 LTRs and flanking AAVS1 homology arms (20). The corresponding CRISPR/Cas9 delivery plasmid PX458_AAVS1 gRNA plasmid encodes Cas9 and a guide RNA (gRNA) targeting the AAVS1 locus (20).

#### Generation of HIV-2 and HTLV-1 LTatCL[M] vectors

HIV-2 and HTLV-1 LTatCL[M] constructs were derived from the HIV-1 LTatCΔL[M] intermediate. To generate this intermediate, a polyadenylation (polyA) fragment was PCR-amplified using CloneAmp HiFi Premix (Takara Bio, 639298) with primers introducing overlaps to the Cerulean cassette and the 3′ HIV-1 LTR, an AgeI site, and an AsiSI repair sequence. The fragment was inserted into AsiSI-linearised HIV-1 LTatCL[M] by In-Fusion cloning (In-Fusion HD Cloning Kit, Takara Bio, 639649), generating HIV-1 LTatC-polyA-L[M]. The 3′ LTR was subsequently excised by AgeI-HF and partial EcoRI-HF digestion, and an adapter containing a unique PmeI site and ends complementary to the linearised vector was generated from annealed oligonucleotides and ligated into the backbone using T7 DNA ligase, generating HIV-1 LTatCΔL[M].

HIV-1 LTatCΔL[M] was digested with AsiSI and AgeI-HF to excise the polyA region and purified. The HIV-2 3′ LTR was amplified from an HIV-2 ST infectious clone (kind gift of Dr Beatrice Hahn and Dr George M. Shaw (40, 41) ) using primers with overlaps to the Cerulean cassette and adapter region and CloneAmp HiFi PCR Premix, and the product was purified. This PCR product was inserted into the linearised vector by In-Fusion cloning (insert:vector molar ratio 5:1) to replace the polyA region and deleted 3′ LTR. The resulting intermediate plasmid was linearised with AscI (NEB, R0558S), partially digested with SalI-HF (NEB, R3138S) for 45 min to remove the 5′ LTR, and gel-purified. The HIV-2 5′ LTR was amplified in the same way using primers with overlaps to the 3′ end of the AAVS1 5′ arm and the 5′ end of the *tat* region, purified, and cloned into this backbone by In-Fusion cloning to yield the HIV-2 LTatCL[M] vector. An analogous strategy was used to generate the HTLV-1 LTatCL[M] construct, using HTLV-1 LTRs amplified from MT-2 cellular DNA carrying integrated HTLV-1 provirus as template.

#### Generation of ΔLTatCΔL[M]

To generate the LTR-deficient construct ΔLTatCΔL[M], HIV-1 LTatCΔL[M] was digested with AscI and subjected to partial digestion with SalI-HF for 45 min to excise the 5′ LTR, followed by gel extraction and purification. The backbone was combined in an In-Fusion reaction (insert:vector molar ratio 5:1) with an EF1α promoter fragment designed with overlaps to the 3′ end of the AAVS1 5′ arm and the 5′ end of the tat region. The elongation factor 1 alpha (EF1α) fragment was amplified using CloneAmp HiFi Premix. The resulting ΔLTatCΔL[M] plasmid was verified by Sanger sequencing.

#### Plasmid propagation and verification

PCR products and restriction fragments were purified, including gel extraction where required, using the NucleoSpin Gel and PCR Clean-up Kit (Macherey-Nagel, 740609.50). All ligation and In-Fusion reactions were transformed into Stellar Competent Cells (Takara Bio, 636766) and plated on LB agar containing 100 µg/ml ampicillin (Sigma-Aldrich, A0166). Single colonies were inoculated into LB broth (Sigma-Aldrich, L3022) with ampicillin and grown overnight at 37°C. Primer design, cloning, and bacterial transformation followed the In-Fusion HD Cloning Kit user manual (Takara Bio). Plasmid DNA was isolated using the NucleoSpin Plasmid Mini kit for mini-preparations (Macherey-Nagel, 740588.50) and the QIAGEN Plasmid Maxi Kit (Qiagen, 12165) for large-scale preparations. Plasmid DNA concentrations were measured by NanoDrop One C microvolume spectrophotometer (Thermo Scientific), and Qubit 4 fluorometer (Invitrogen). All final constructs were confirmed by Sanger sequencing and subsequently by NGS. HPLC-purified primers and oligonucleotides were ordered from Microsynth AG and dissolved in Tris buffer. Primer sequences and amplification conditions are listed in Supplementary Table S1 to S6.

### Nucleofection of K562 cells

Four LTatCL[M] vectors (HIV-1 LTatCL[M], HIV-2 LTatCL[M], HTLV-1 LTatCL[M], and ΔLTatCΔL[M]) were linearised with NsiI-HF (New England Biolabs, R3127S) before nucleofection. For each condition, 3 × 10⁶ K562 cells were nucleofected with 2.5 µg of the respective linearised dual-fluorophore vector and 2.5 µg of PX458_AAVS1 gRNA plasmid using an Amaxa Nucleofector II (Lonza) and Nucleofector 2b Kit V (Lonza, VCA-1003) with programme T-016, following the manufacturer’s protocol for K562 cells. Nucleofections were performed in quadruplicate per vector, and nucleofected cells were transferred to 6-well plates at 3 × 10⁶ cells in 2.5 ml complete medium per well.

### Flow cytometry and cell sorting

Nucleofected K562 cells were analysed on a CytoFLEX S flow cytometer (Beckman Coulter) to visualise and quantify Cerulean and mCherry fluorescence. Cell viability was assessed by staining with Zombie NIR dye (Zombie NIR Fixable Viability Kit, BioLegend, 423105) diluted 1:1000 in PBS, according to the manufacturer’s instructions. In brief, 1 × 10⁶ cells were resuspended in 100 µl of diluted dye, incubated for 20 min at room temperature in the dark, washed at least twice with PBS, and subsequently processed in FACS buffer for acquisition. Zombie NIR staining was performed only for analytical flow cytometry on the CytoFLEX S. On the CytoFLEX S, Cerulean and mCherry fluorescence were visualised using bivariate plots with quadrant gates to define double-positive Cerulean⁺/mCherry⁺, single mCherry⁺ and single Cerulean⁺ populations. Gating strategies were established using non-transfected controls and kept similar across experiments.

DP, single mCherry⁺ and single Cerulean⁺ populations were isolated by fluorescence-activated cell sorting (FACS) at the Cytometry Facility of the University of Zurich using a BD FACSAria III (BD Biosciences). For FACS sorting, sequential gates were applied to select cells, singlets and then polygon-gated Cerulean/mCherry populations; more permissive gates were used during first round of sorting to enrich populations for expansion, and stricter gates during second round of sorting to obtain highly pure populations for downstream analysis. Nucleofected K562 populations were expanded after the initial sorting and subjected to a second round of sorting; prior to the second sort, single mCherry⁺ cells were treated for 24 h with 10 ng/ml TNFα and 1 µM SAHA to reactivate transcriptionally silent reporter integrations (a proxy for latent proviruses). Flow cytometry data, including gating, were analysed using FlowJo v10 (BD Biosciences).

### Pulsed-field gel electrophoresis

PFGE was used to enrich for high-molecular-weight genomic DNA harbouring integrated copies of the model vector and to remove episomal or short DNA species or unintegrated vector. PFGE runs were carried out on a BluePippin instrument (Sage Science) as was previously described for selective recovery of long genomic fragments (22, 42) , using gel cassettes and reagents supplied by the manufacturer (kit BLF7510). Samples were processed according to the manufacturer’s protocol with the cassette setting “0.75% DF Marker S1 high-pass 15–20 kb” to retain fragments larger than 15 kb and deplete shorter or sheared DNA fragments.

Genomic DNA was extracted using the DNeasy Blood and Tissue Kit (Qiagen, 69506), initially quantified with a NanoDrop, concentrated by evaporation, and re-quantified using a Qubit. Between 500 ng and 5 µg DNA in a maximum volume of 30 µl was loaded per lane, in line with the instrument recommendation of ≤5 µg DNA per lane. Eluted DNA fractions were collected and quantified again with Qubit.

### Mapping of large internal deletions

PFGE-purified DNA from DP, non-inducible single mCherry⁺, reactivated DP and single Cerulean⁺ populations was used for amplification to map deletions within the Cerulean and mCherry cassettes. Genomic DNA concentrations were adjusted such that each PCR reaction contained DNA in the low-nanogram range, corresponding approximately to a single or few vector copies per reaction. For each sample, multiple independent PCRs (typically ≥16) were set up using different template dilutions (for example 1:5 or 1:10) to optimise conditions for obtaining discrete bands suitable for sequencing.

Each 25 µl PCR contained 5 µl 5x LongAmp Taq Reaction Buffer (New England Biolabs, M0534L), 0.75 µl 10 mM dNTP mix (Thermo Fisher Scientific, R0181), 1.25 µl 10 µM forward primer, 1.25 µl 10 µM reverse primer, 1 µl LongAmp Hot Start Taq DNA Polymerase (NEB, M0534L), and nuclease-free water to volume; 1 µl of diluted PFGE-purified DNA was added as template.

Amplicons were resolved by agarose gel electrophoresis to assess product size, and reactions yielding single or very few discrete bands were purified using the NucleoSpin Gel and PCR Clean-up Kit (Macherey-Nagel, 740609.50) for downstream next-generation sequencing. Primer sequences and cycling conditions are provided in Supplementary table S1 and S7 respectively.

### Next-generation sequencing (NGS)

To characterise deletions in the model vector, we performed NGS on PCR-purified amplicons generated from PFGE-purified genomic DNA from DP, single mCherry⁺, and single Cerulean⁺ populations. DNA concentration was determined with a Qubit fluorometer and samples were adjusted to 0.2 ng/µl before library preparation. Libraries were generated using the Nextera XT DNA Library Prep Kit (Illumina, FC-131-1096) together with the Nextera XT Index Kit v2 Set A (Illumina, FC-131-2001). Sequencing was carried out on an Illumina MiSeq instrument with the MiSeq Reagent Kit v3, 150-cycle configuration (Illumina, MS-102-3001), following the manufacturer’s instructions and including a 1% PhiX control spike-in.

All plasmid constructs used in this study were sequence verified by NGS using the same workflow. Raw sequencing data were retrieved from the openBIS system (University of Zurich) and processed with CLC Genomics Workbench 25 (Qiagen).

## Supporting information

Supplementary file 1 with table S1 to S7

## Data availability

All data underlying the results presented in this study are included in the article and its Supporting Information files. Raw next-generation sequencing data generated in this study have been deposited in the European Nucleotide Archive (ENA) under study accession PRJEB128069 and will be made publicly available upon publication. Additional underlying data can be obtained from the corresponding author upon reasonable request.

## Abbreviations

gRNA: Guide RNA
HIV-1: Human immunodeficiency virus type 1
HIV-2: Human immunodeficiency virus type
HTLV-1: Human T-lymophotropic virus
LID: Large internal deletion
DP: Double-positive
MMEJ: Microhomology mediated end joining
LTR: Long terminal repeat
PFGE: Pulsed-field gel electrophoresis
TNFα: Tumor necrosis factor
SAHA: Suberoylanilide hydroxamic acid
AAVS1: Adeno-associated virus integration site 1
LRA: Latency reversal agent
HDAC: Histone deacetylase

## Acknowledgments

We thank Manuel Schulthess, Mario Wicker, and Priyanikha Jayakumar (Flow Cytometry Facility, University of Zurich [UZH], Zurich, Switzerland) for bulk cell-sorting services; Gabriela Ziltener (Institute of Medical Virology, UZH, Zurich, Switzerland) for next-generation sequencing; Dr Anne Inderbitzin and Dr Tom Loosli for providing the HIV-1 LTatCL[M] plasmid and for assistance in generating the four final model vectors used in this study; and Aria Maya Minder Pfyl (Genetic Diversity Centre, ETH Zurich, Zurich, Switzerland) for access to the BluePippin device. The following reagent was obtained through the NIH HIV Reagent Program, Division of AIDS, NIAID, NIH: Human Immunodeficiency Virus Type 2 (HIV-2) ST Infectious Molecular Clone, ARP-12444, contributed by Dr. Beatrice Hahn and Dr. George Shaw.

The manuscript’s text was edited for grammar and clarity using an AI-based language tool (Perplexity); all scientific content and interpretations are the authors’ own.

This study was supported by the Swiss National Science Foundation (SNF) (grant number 310030_204404 to Karin J. Metzner (K.J.M).). The funder had no role in the design of the study; in the collection, analyses, or interpretation of data; in the writing of the manuscript; or in the decision to publish the results.

Within the last 5 years, K.J.M. has received advisory board honoraria from ViiV, and the University of Zurich received research grants from Gilead Sciences and Novartis for studies for which Dr. Metzner serves as principal investigator.

## Author Contributions

- Sagnik Bhattacharjee conceived and designed the experiments, performed the experiments, analysed the data, prepared figures and/or tables, authored or reviewed drafts of the paper, and approved the final draft.
- Koleta Michalek performed the experiments, analysed the data, reviewed drafts of the paper, and approved the final draft.
- Karin J. Metzner conceived and designed the experiments, analysed the data, authored or reviewed drafts of the paper, and approved the final draft.

## Supplementary information

Supplementary File 1. Supplementary Table S1-S7 containing primer sequences and cycling conditions.

Supplementary Figure S1. Enrichment of defined Cerulean⁺ and/or mCherry⁺ subpopulations for analysis of large internal deletions in HIV-2 LTatCL[M].

Supplementary Figure S2. Enrichment of defined Cerulean⁺ and/or mCherry⁺ subpopulations for analysis of large internal deletions in HTLV-1 LTatCL[M].

Supplementary Figure S3. Enrichment of defined Cerulean⁺ and/or mCherry⁺ subpopulations for analysis of large internal deletions in ΔLTatCΔL[M].

Supplementary Figure S4. Large internal deletions in single Cerulean⁺ populations

Supplementary Figure S5. Intact and deleted vector genomes in double-positive and reactivated double-positive populations.

