## Supplementary figures S1 to S5 for "Long terminal repeats influence deletion propensity and genome instability in proviruses across major retroviruses"

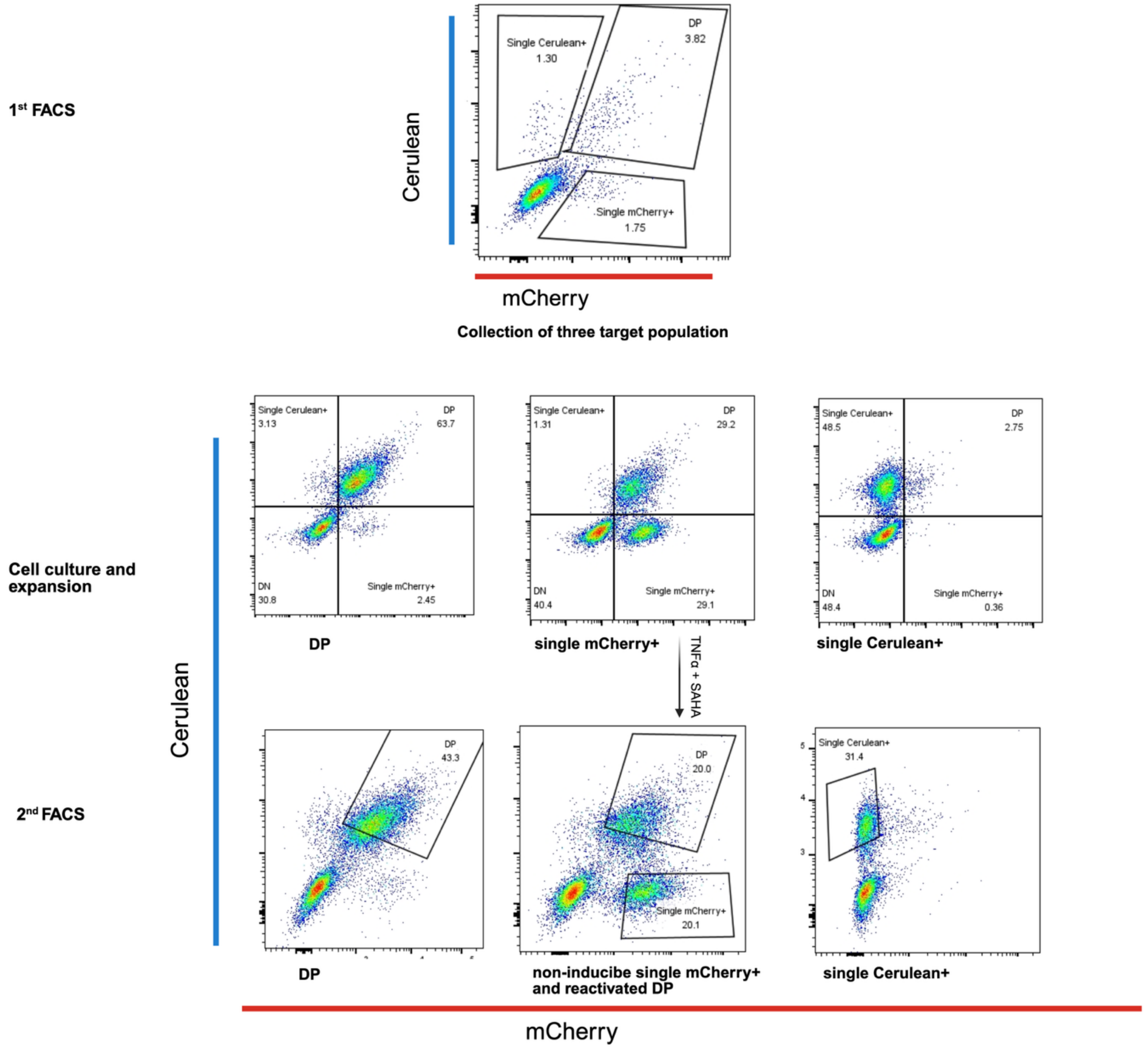


**Figure S1. Enrichment of defined Cerulean$^+$ and/or mCherry$^+$ subpopulations for analysis of large internal deletions in HIV-2 LTatCL[M].**

Representative flow cytometry plots for HIV-2 LTatCL[M]. Top row: gating at first sort (day 6 after nucleofection) to collect double-positive (DP), single Cerulean$^+$, and single mCherry$^+$ populations. Middle row: analysis of the expanded populations at day 16. Bottom row: gating at the second sort (day 18) to collect DP from the DP culture, non-inducible single mCherry$^+$ and reactivated DP from LRA-treated single mCherry$^+$ cultures, and single Cerulean$^+$ from single Cerulean$^+$ cultures. Flow cytometry data from both analytical runs and sorting (CytoFLEX and BD FACSAria) were reanalysed in FlowJo to apply consistent gates, axis orientation, and labelling across experiments; plots were arranged and refined in BioRender without altering underlying event distributions.

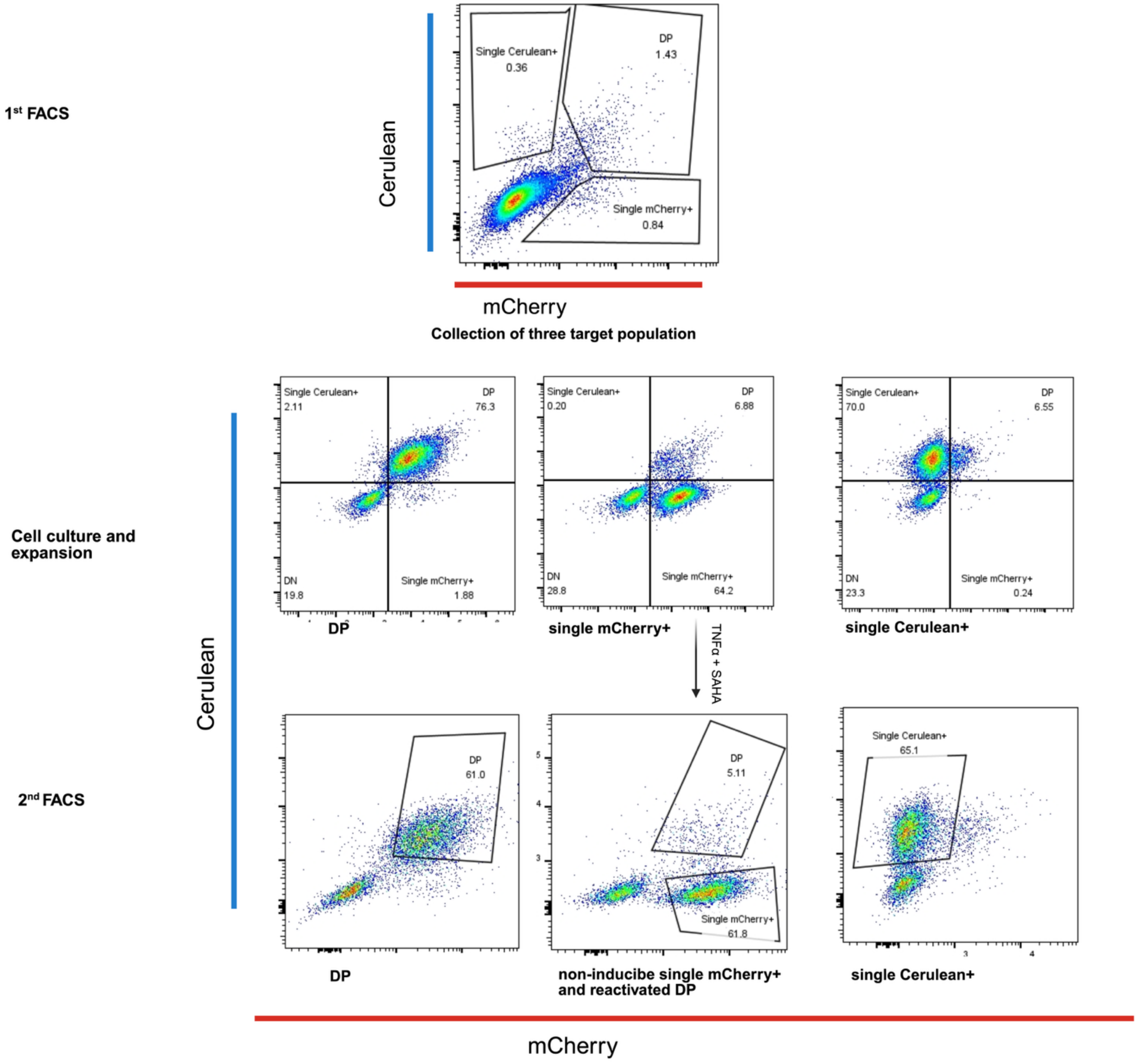


**Figure S2. Enrichment of defined Cerulean$^{+}$ and/or mCherry$^{+}$ subpopulations for analysis of large internal deletions in HTLV-1 LTatCL[M].**

Representative flow cytometry plots for HTLV-1 LTatCL[M]. Top row: gating at first sort (day 8 after nucleofection) to collect double-positive (DP), single Cerulean$^+$, and single mCherry$^+$ populations. Middle row: analysis of the expanded populations at day 20. Bottom row: gating at the second sort (day 24) to collect DP from the DP culture, non-inducible single mCherry$^+$ and reactivated DP from LRA-treated single mCherry$^+$ cultures, and single Cerulean$^+$ from single Cerulean$^+$ cultures. Flow cytometry data from both analytical runs and sorting (CytoFLEX and BD FACSAria) were reanalysed in FlowJo to apply consistent gates, axis orientation, and labelling across experiments; plots were arranged and refined in BioRender without altering underlying event distributions.

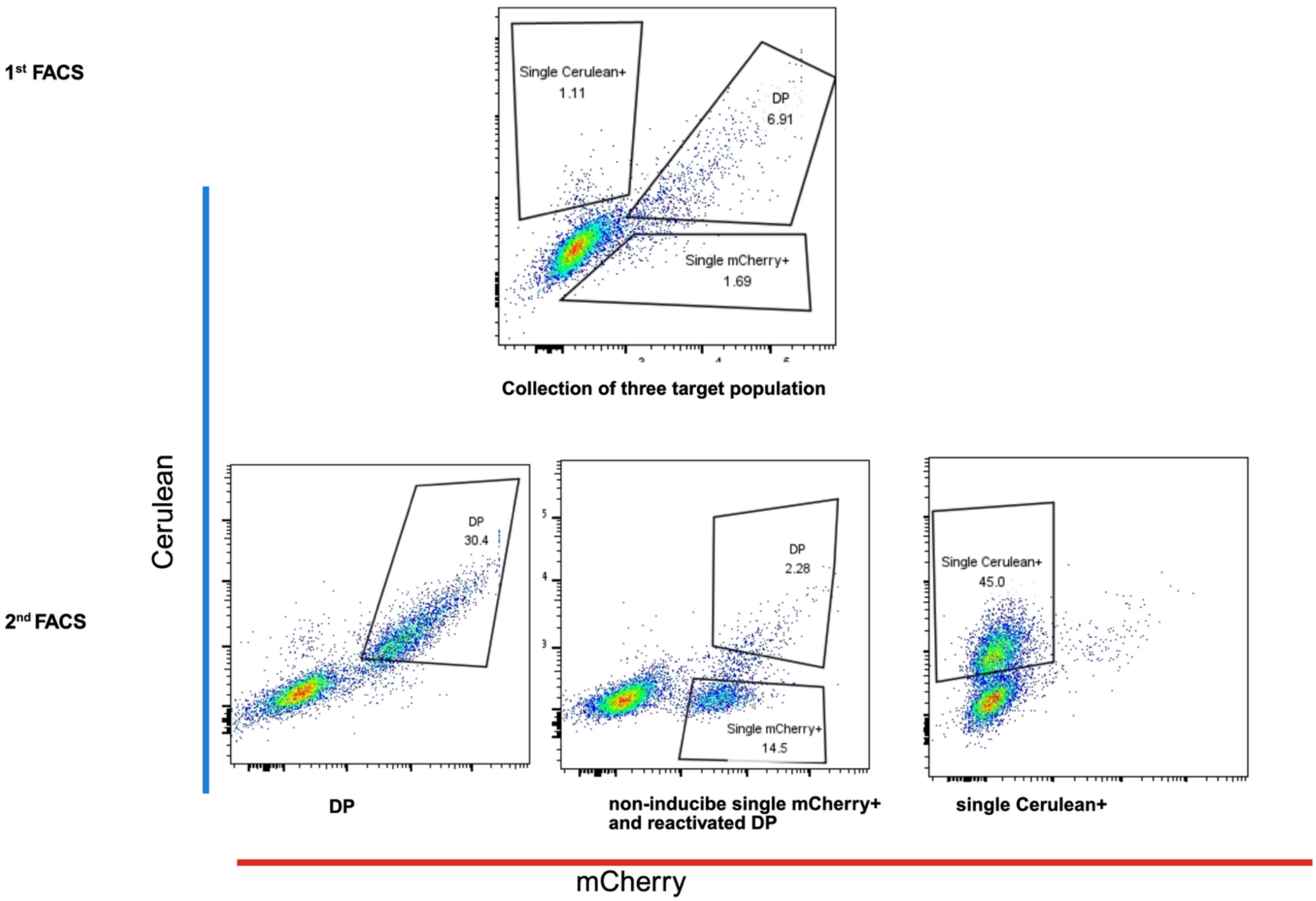


**Figure S3. Enrichment of defined Cerulean$^+$ and/or mCherry$^+$ subpopulations for analysis of large internal deletions in ΔLTatCΔL[M].**

Representative flow cytometry plots for ΔLTatCΔL[M]. Top row: gating at first sort (day 6 after nucleofection) to collect double-positive (DP), single Cerulean$^+$, and single mCherry$^+$ populations. Bottom row: gating at the second sort (day 18) to collect DP from the DP culture, non-inducible single mCherry$^+$ and reactivated DP from LRA-treated single mCherry$^+$ cultures, and single

Cerulean$^+$ from single Cerulean$^+$ cultures. Flow cytometry data from both analytical runs and sorting (CytoFLEX and BD FACSAria) were reanalysed in FlowJo to apply consistent gates, axis orientation, and labelling across experiments; plots were arranged and refined in BioRender without altering underlying event distributions.

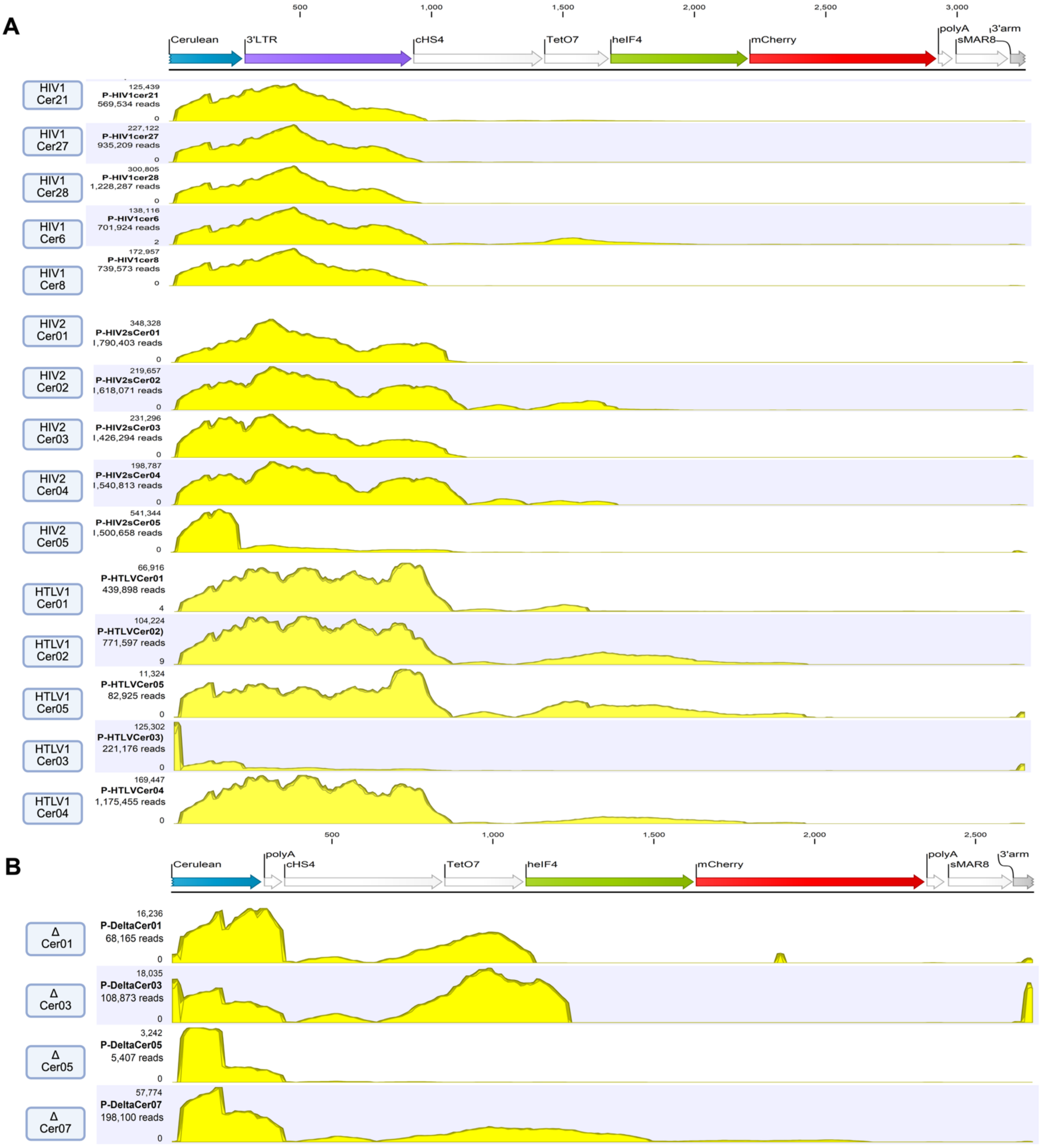

A
500
1,000
1,500
2,000
2,500
3,000
Cerulean
3'LTR
cHS4
TetO7
heIF4
mCherry
polyA
sMAR8
3'arm
HIV1 Cer21
125,439
P-HIV1cer21
569,534 reads
HIV1 Cer27
227,122
P-HIV1cer27
935,209 reads
HIV1 Cer28
300,805
P-HIV1cer28
1,228,287 reads
HIV1 Cer6
138,116
P-HIV1cer6
701,924 reads
HIV1 Cer8
172,957
P-HIV1cer8
739,573 reads
HIV2 Cer01
348,328
P-HIV2sCer01
1,790,403 reads
HIV2 Cer02
219,657
P-HIV2sCer02
1,618,071 reads
HIV2 Cer03
231,296
P-HIV2sCer03
1,426,294 reads
HIV2 Cer04
198,787
P-HIV2sCer04
1,540,813 reads
HIV2 Cer05
541,344
P-HIV2sCer05
1,500,658 reads
HTLV1 Cer01
66,916
P-HTLVCer01
439,898 reads
HTLV1 Cer02
104,224
P-HTLVCer02)
771,597 reads
HTLV1 Cer05
11,324
P-HTLVCer05
82,925 reads
HTLV1 Cer03
125,302
P-HTLVCer03)
221,176 reads
HTLV1 Cer04
169,447
P-HTLVCer04
1,175,455 reads
B
500
1,000
1,500
2,000
2,500
Cerulean
polyA
cHS4
TetO7
heIF4
mCherry
polyA
sMAR8
3'arm
Δ Cer01
16,236
P-DeltaCer01
68,165 reads
Δ Cer03
18,035
P-DeltaCer03
108,873 reads
Δ Cer05
3,242
P-DeltaCer05
5,407 reads
Δ Cer07
57,774
P-DeltaCer07
198,100 reads

**Figure S4. Large internal deletions in single Cerulean+ populations.**

Coverage plots of all amplicon sequences per vector showing deletions within the mCherry cassette in single Cerulean+ populations for (A) HIV-1 LTatCL[M], HIV-2 LTatCL[M], and HTLV-1 LTatCL[M], and (B) the LTR-deficient ΔLTatCΔL[M] construct. At the top, a schematic of the corresponding reference mCherry-containing region is shown. Yellow peaks indicate regions with read coverage (intact sequence), whereas flat gaps between or after peaks indicate internal deletions. Individual amplicon identifiers (left) correspond to specific PCR products and sequencing reactions and match the sequence names used for data deposition. Coverage plots were exported from CLC Genomics Workbench after mapping reads to the corresponding reference constructs and assembled in BioRender.

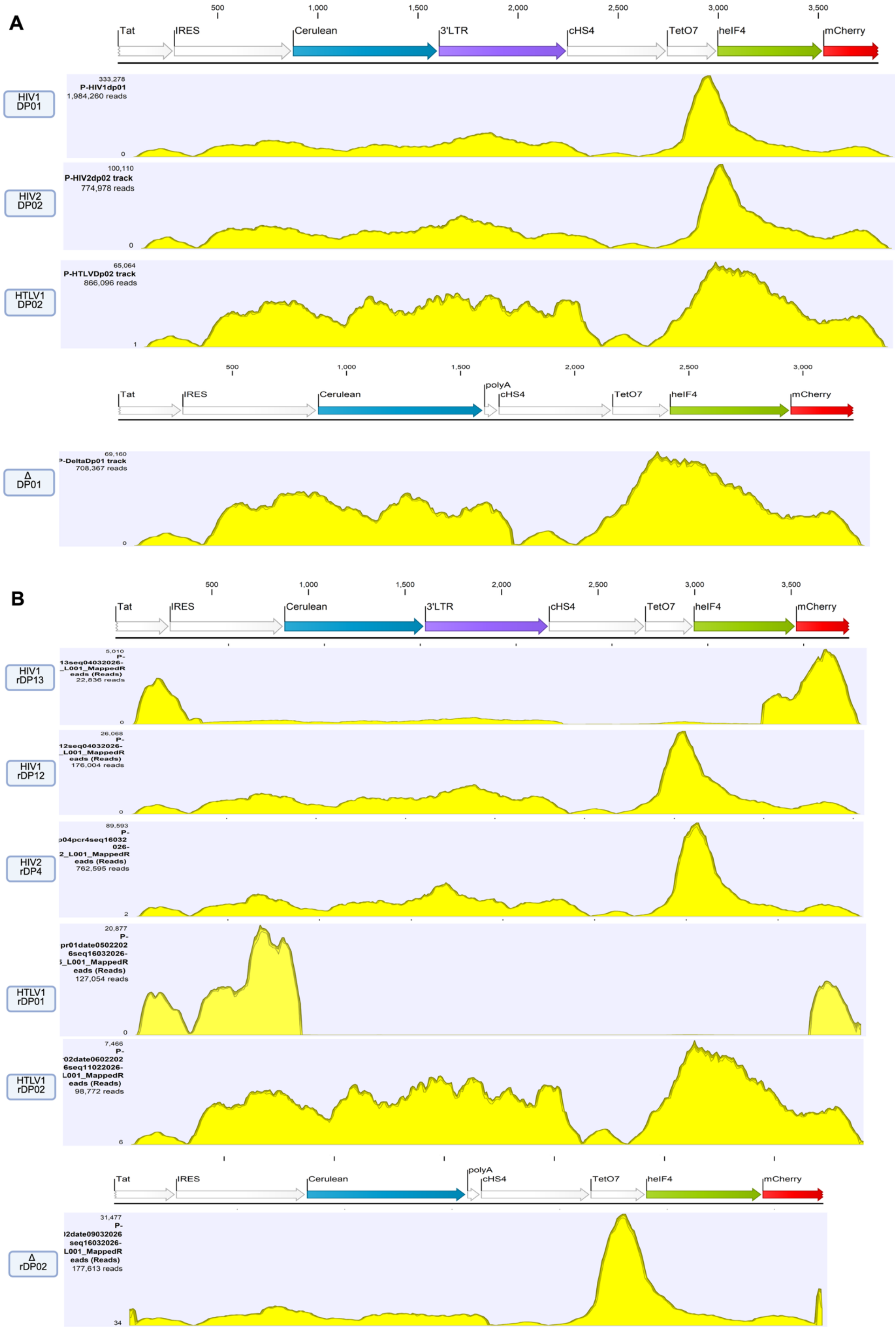
A
500
1,000
1,500
2,000
2,500
3,000
3,500
Tat
IRES
Cerulean
3'LTR
cHS4
TetO7
heIF4
mCherry
HIV1 DP01
333,278
P-HIV1dp01
1,984,260 reads
0
HIV2 DP02
100,110
P-HIV2dp02 track
774,978 reads
0
HTLV1 DP02
65,064
P-HTLVDp02 track
866,096 reads
1
polyA
Δ DP01
69,160
708,367 reads
0
B
HIV1 rDP13
5,010
22,836 reads
0
HIV1 rDP12
26,068
176,004 reads
0
HIV2 rDP4
89,593
762,595 reads
2
HTLV1 rDP01
20,877
127,054 reads
0
HTLV1 rDP02
7,466
98,772 reads
6
Δ rDP02
31,477
177,613 reads
34

**Figure S5. Intact and deleted vector genomes in double-positive and reactivated double-positive populations.**

A) Representative coverage plots of one amplicon sequence per vector from double-positive (DP) populations for HIV-1 LTatCL[M], HIV-2 LTatCL[M], HTLV-1 LTatCL[M], and LTR-deficient ΔLTatCΔL[M] construct, showing intact integrated vectors. Corresponding reference schematics are shown at the top.

B) Representative coverage plots of reactivated double-positive (rDP) amplicon sequences for HIV-1 LTatCL[M], HIV-2 LTatCL[M], HTLV-1 LTatCL[M], and the LTR-deficient ΔLTatCΔL[M] construct, with corresponding reference schematics shown at the top. For HIV-1 LTatCL[M] and HTLV-1 LTatCL[M], one intact and one internally deleted rDP sequence are displayed to illustrate that rare internal deletions can occur without loss of reporter expression. Intact rDP sequences are shown for HIV-2 LTatCL[M] and ΔLTatCΔL[M], for which no deletions were observed. Yellow peaks indicate regions with read coverage (intact sequence), whereas flat gaps between or after peaks indicate internal deletions. Individual amplicon identifiers (left) correspond to specific PCR products and sequencing reactions and match the sequence names used for data deposition. Coverage plots were exported from CLC Genomics Workbench after mapping reads to the corresponding reference constructs and assembled in BioRender.
