## Supplementary file 1 with table S1 to S7 for "Long terminal repeats influence deletion propensity and genome instability in proviruses across major retroviruses"

**Table S1. Primers for CloneAmp HiFi PCR and Infusion Cloning and to map deletions:**

| **Name** | **Sequences(5'to3')** | **Purpose** |
| --- | --- | --- |
| polyAfrag_0925_Fw | CTCGGCATGGACGAGCTGTACAAGTAAGCGATCGCAATAAAAGATCTTTATTTTCATTAG | In-Fusion cloning amplification of polyA fragment into HIV-1 LTatCL[M] |
| polyAfrag_0925_Rv | TTTTTGGGACCAAATTAGCCCTTCCAGTACCGGTGCTCTAGACACACAAAAAACCAACAC |  |
| Adapter_191008_fw | CCGGTCTACCTGGGTTTAAACGAGGTGACG | Oligonucleotides for adapter ligation with restriction site for PmeI |
| Adapter_191008_rv | AATTCGTCACCTCGTTTAAACCCAGGTAGA |  |
| HIV 2 _5' LTR_Ltat_cloning_for | AGGGACATTAGGCGCGCCTGGAAGGGCTGTATTACAGTG | In-Fusion cloning amplification of 5' LTR of HIV-2 into LTatCΔL[M] replacing HIV- 1 5' LTR |
| HIV 2 _5' LTR_Ltat_cloning_rev | GCTCCATTTCGTCGACGCTAGGGATTTTCCTGCCTTG |  |
| HIV 2 _3' LTR_Ltat_cloning_for | CTGTACAAGTAAGCGATCGCTGGAAGGGCTGTATTACAGTG | In-Fusion cloning amplification of 3' LTR of HIV-2 into LTatCΔL[M] replacing poly A and deleted HIV-1 3' LTR |
| HIV 2 _3' LTR_Ltat_cloning_rev | ACCCAGGTAGACCGGTGCTAGGGATTTTCCTGCCTTG |  |
| HTLV 1 _5' LTR_Ltat_cloning_for | AGGGACATTAGGCGCGCCTGACAATGACCATGAGCCCCAAAT | In-Fusion cloning amplification of 5' LTR of HTLV-1 into LTatCΔL[M] replacing HIV - 1 5' LTR |
| HTLV 1 _5' LTR_Ltat_cloning_rev | GCTCCATTTCGTCGACTGTGTACTAAGTTTCTCTCCTGGAG |  |
| HTLV 1 _3' LTR_Ltat_cloning_for | CTGTACAAGTAAGCGATCGCTGACAATGACCATGAGCCCCAAAT | In-Fusion cloning amplification of 3' LTR of HTLV-1 into LTatCΔL[M] replacing poly A and deleted HIV-1 3' LTR |
| HTLV 1 _3' LTR_Ltat_cloning_rev | ACCCAGGTAGACCGGTGTGTACTAAGTTTCTCTCCTGGAG |  |
| Elf1a_191010_FW | CACAGTGGGGCCACTAGGGACATTAGCGTACGGAGTAATTCATACAAAAGGACTCGCCCC | In-Fusion Cloning amplification of eF1α fragment into LTatCΔL[M] replacing HIV - 1 5' LTR |
| Elf1a_191010_RV | GGATCTACTGGCTCCATTTCGTCGAGTTAACTCACGACACCTGAAATGGAAGAAAAAAAC |  |
| Cerulean full-length tat fw | GGAAGCATCCAGGAAGTCAGC | Amplification of Cerulean cassette from tat to the start of mCherry to map deletions. |
| Cerulean full length mch rev | GAAGGACAGCTTCAAGTAGTCG |  |
| mCherry full length cer fw | CAGCGACAACGTCTATATCACC | Amplification of the mCherry cassette (middle of Cerulean to 3′ AAVS1 arm) to map deletions. |
| AAVS LtatCLM rev | CCAATATCAGGAGACTAGGAAGG |  |

**Table S2. Amplification of polyA fragment:**

| 98°C | 10 sec |  |
| --- | --- | --- |
| 98°C | 10 sec | 35x |
| 57°C | 15 sec |  |
| 72°C | 5 sec |  |
| 72°C | 5 min |  |
| 10°C | hold |  |

The DNA insert (10 ng) was amplified using the CloneAmp HiFi PCR Premix (Takara Bio) according to manufacturer’s instructions and primers (0.5 μM)

**Table S3. Cycling conditions for annealing of adapters**

| 37°C | 30 min Phosphorylation |
| --- | --- |
| 95°C | 5 min Denaturation |
| 25°C | -0.1°C/min Ligation ramp |

**Table S4. Amplification of HIV-2 LTR**

| 95°C | 1 min |  |
| --- | --- | --- |
| 98°C | 10 sec |  |
| 55°C | 15 sec | 35x |
| 72°C | 10 sec |  |
| 72°C | 7 min |  |
| 4°C | hold |  |

The DNA insert (HIV-2 LTR from 1 ng of HIV-2 ST infectious clone) was amplified using the CloneAmp HiFi PCR Premix (Takara Bio) according to manufacturer’s instructions and primers (0.3 μM)

**Table S5. Amplification of HTLV-1 LTR**

| 95°C | 1 min |  |
| --- | --- | --- |
| 98°C | 10 sec |  |
| 55°C | 15 sec | 35x |
| 72°C | 30 sec |  |
| 72°C | 7 min |  |
| 4°C | hold |  |

The DNA insert (HTLV-1 LTR from 100 ng of MT-2 cellular DNA) was amplified using the CloneAmp HiFi PCR Premix (Takara Bio) according to manufacturer’s instructions and primers (0.3 μM)

**Table S6. Amplification of eF1α**

| 98°C | 10 sec |  |
| --- | --- | --- |
| 98°C | 10 sec | 35x |
| 57°C | 15 sec |  |
| 72°C | 15 sec |  |
| 72°C | 5 min |  |
| 10°C | hold |  |

The DNA insert (eF1α from 10 ng of the plasmid) was amplified using the CloneAmp HiFi PCR Premix (Takara Bio) according to manufacturer’s instructions and primers (0.5 μM)

**Table S7. PCR amplification of Cerulean and mCherry cassette to map deletions, if any**

| 94°C | 2min |  |
| --- | --- | --- |
| 94°C | 30s | 35 cycles |
| 58°C | 30s |  |
| 65°C | 5min |  |
| 65°C | 10min |  |
| 10°C | hold |  |
